# Co-localization of Conditional eQTL and GWAS Signatures in Schizophrenia

**DOI:** 10.1101/129429

**Authors:** Amanda Dobbyn, Laura M. Huckins, James Boocock, Laura G. Sloofman, Benjamin S. Glicksberg, Claudia Giambartolomei, Gabriel Hoffman, Thanneer Perumal, Kiran Girdhar, Yan Jiang, Douglas M. Ruderfer, Robin S. Kramer, Dalila Pinto, the CommonMind Consortium, Schahram Akbarian, Panos Roussos, Enrico Domenici, Bernie Devlin, Pamela Sklar, Eli A. Stahl, Solveig K. Sieberts

## Abstract

Causal genes and variants within genome-wide association study (GWAS) loci can be identified by integrating GWAS statistics with expression quantitative trait loci (eQTL) and determining which SNPs underlie both GWAS and eQTL signals. Most analyses, however, consider only the marginal eQTL signal, rather than dissecting this signal into multiple independent eQTL for each gene. Here we show that analyzing conditional eQTL signatures, which could be important under specific cellular or temporal contexts, leads to improved fine mapping of GWAS associations. Using genotypes and gene expression levels from post-mortem human brain samples (N=467) reported by the CommonMind Consortium (CMC), we find that conditional eQTL are widespread; 63% of genes with primary eQTL also have conditional eQTL. In addition, genomic features associated with conditional eQTL are consistent with context specific (i.e. tissue, cell type, or developmental time point specific) regulation of gene expression. Integrating the Psychiatric Genomics Consortium schizophrenia (SCZ) GWAS and CMC conditional eQTL data reveals forty loci with strong evidence for co-localization (posterior probability >0.8), including six loci with co-localization of conditional eQTL. Our co-localization analyses support previously reported genes and identify novel genes for schizophrenia risk, and provide specific hypotheses for their functional follow-up.

## INTRODUCTION

Significant advances in understanding the genetic architecture of schizophrenia have occurred over the last ten years. However, for common variants identified in genome-wide association studies (GWAS), the success in locus identification is not yet matched by an understanding of their underlying basic mechanism or effect on pathophysiology. Expression quantitative trait loci (eQTL), which are responsible for a significant proportion of variation in gene expression, could serve as a link between the numerous non-coding genetic associations that have been identified in GWAS and susceptibility to common diseases directly through their association with gene expression regulation.^1-4^ Indeed, results from eQTL mapping studies have been successfully utilized to identify genes and causal variants from GWAS for various complex phenotypes, including asthma, body mass index, celiac disease, and Crohn’s disease.^5-8^

Studies integrating eQTL and GWAS data have almost exclusively used marginal association statistics which typically represent the primary, or most significant, eQTL signal when assessing co-localization with GWAS, ignoring other SNPs that affect expression independently of the primary eQTL for a given gene. However, recent findings indicating that conditionally independent eQTL are widespread^9-11^ motivate examination of the extent to which considering conditional eQTL may provide additional power to identify likely causal genes in a GWAS locus. Recent reports provide evidence that conditional eQTL are less frequently shared across tissues than primary eQTL^9^ and, like tissue and cell type specific eQTL, are often found more distally to the genes they regulate.^9; 12; 13^ These lines of evidence suggest that conditionally independent eQTL may contribute to tissue- or other context-specific gene regulation (e.g. specific to a particular cell type, developmental stage, or stimulation condition).

Here, we leveraged genotype and dorsolateral prefrontal cortex (DLPFC) expression data provided by the CommonMind Consortium (CMC) to elucidate the role of conditional eQTL in the etiology of schizophrenia (SCZ). Currently comprising the largest existing postmortem brain genomic resource at nearly 600 samples, the CMC is generating and making publicly available an unprecedented array of functional genomic data, including gene expression (RNA-sequencing), histone modification (chromatin immunoprecipitation, ChIP-seq), and SNP genotypes, from individuals with psychiatric disorders as well as unaffected controls.^14^ We utilized SNP dosage and RNA-sequencing (RNA-seq) data from the CMC to identify primary and conditionally independent eQTL. We then characterized the resulting eQTL on various genomic attributes including distance to transcription start site, and their genes’ specificity across tissues, cell-types, and developmental periods. In addition, we quantified enrichment of primary and conditional eQTL in promoter and enhancer functional genomic elements inferred from epigenomic data. Finally, we isolated each independent eQTL signal by conducting a series of “all-but-one” conditional analyses for genes with multiple independent eQTL, and assessed the overlap between all eQTL association signals and the SCZ GWAS signals.

## MATERIAL AND METHODS

### CommonMind Consortium Data

We used pre-QC’ed genotype and expression data made available from the CommonMind Consortium, and detailed information on quality control, data adjustment and normalization procedures can be found in Fromer et. al.^14^ Briefly, samples were genotyped at 958,178 markers using the Illumina Infinium HumanOmniExpressExome array, and markers were removed on the basis of having no alternate alleles, having a genotyping call rate 0.98, or a Hardy-Weinberg P-value < 5×10^-5^. After phasing and imputation using the 1000 Genomes Phase 1 integrated reference then filtering out variants with INFO < 0.8 or MAF < 0.05, the total number of markers included in the analysis increased to approximately 6.4 million. Gene expression was assayed via RNA-seq using 100 base pair paired end reads, and mapped to human Ensembl gene reference (v70) using TopHat version 2.0.9 and Bowtie version 2.1.0. After discarding genes with less than 1 CPM (counts per million) in at least 50% of the samples, RNA-seq data for a total of 16,423 Ensembl genes were considered for analysis. The expression data was voom-adjusted for both known covariates (RIN, library batch, institution, diagnosis, post-mortem interval, and sex) and surrogate variable analysis (SVA) identified surrogate variables. After the removal of individuals that did not pass RNA sample QC (including but not limited to: having RIN < 5.5, having less than 50 million total reads or more than 5% of reads aligning to rRNA, having any discordance between genotyping and RNA-seq data, and having RNA outlier status or evidence for contamination), and retaining only genetically-identified European-ancestry individuals, a total of 467 samples were used for downstream analyses. These 467 individuals comprised 209 SCZ cases, 52 AFF (Bipolar, Major depressive disorder, or Mood disorder, unspecified) cases, and 206 controls.

### eQTL Identification

To identify primary and conditional cis-eQTL, we a conducted forward stepwise conditional analysis implemented in MatrixEQTL^15^ using genotype data at 6.4 million markers and RNA-seq data for 16,423 genes. For each gene with at least one cis-eQTL (gene ± 1 Mb) association at a 5% false discovery rate (FDR), the most significant SNP was added as a covariate in order to identify additional independent associations. This procedure was repeated iteratively until no further FDR significant eQTL were identified. We used a linear regression model, adjusting for diagnosis and five ancestry covariates inferred by GemTools. Following eQTL identification, only autosomal eQTL were retained for downstream analyses.

### Replication in Independent Datasets

Replication was performed in HBCC microarray cohort (dbGaP ID phs000979, see Web Resources) and in the ROSMAP^16^ RNA-seq cohort by fitting the stepwise regression models identified in the CMC data. For cases in which a marker was unavailable in the replication cohort, all models including that marker (i.e. for that eQTL and higher-order eQTL conditional on it, for a given gene) were omitted from replication.

Data from the HBCC cohort was QC’ed and normalized as described in Fromer et al.^14^ DLPFC tissue was profiled on the Illumina HumanHT-12_V4 Beadchips and normalized in an analogous manner to the CMC data. Genotypes were obtained using the HumanHap650Yv3 or Human1MDuov3 chips and imputed to 1000 Genomes Phase 1. Replication of the eQTL models was performed on 279 genetically inferred Caucasian samples (76 controls, 72 SCZ, 43 BP, 88 MDD) adjusting for diagnosis and five ancestry components.

ROSMAP data were obtained from the AMP-AD Knowledge Portal (see Web Resources). Quantile normalized FPKM expression values were adjusted for Age of Death, RIN, PMI, and hidden confounders from SVA, conditional on diagnosis. Only genes with FPKM > 0 in > 50 samples were considered in analyses. QC’ed genotypes were also obtained from the AMP-AD Knowledge Portal and imputed to the Haplotype Reference Consortium (v1.1)^17^ reference panel via the Michigan Imputation Server.^18^ Only markers with imputation quality score R^2^ ≥ 0.7 were considered in the replication analysis. GemTools was used to infer ancestry components as for CMC above. After QC, 494 samples were used for eQTL replication in a linear regression model that also adjusted for diagnosis (Alzheimer’s disease, mild cognitive impairment, no cognitive impairment and other) and four ancestry components.

### Modeling Number of eQTL per Gene on Genomic Features

We considered three genomic features (gene length, number of LD blocks in the cis-region, and genic constraint score) for our modeling analyses. Gene lengths were calculated using Ensembl gene locations. We obtained LD blocks from the LDetect Bitbucket site to tally the number of LD blocks overlapping each gene’s cis-region (gene ± 1Mb). We obtained Loss-of-Function-based genic constraint scores from the Exome Aggregation Consortium (ExAC). A negative-binomial generalized linear regression model was used to model the number of eQTL per gene based on the above variables; results were qualitatively the same using linear regression of Box-Cox transformed eQTL numbers. Backward-forward stepwise regression using the full model with interaction terms for these three variables was used to determine the relationship between genomic conditions and eQTL number. These analyses were implemented in R.

Cis-heritability of gene expression was estimated using the same CMC data used for eQTL detection, using all markers in the cis-region using GCTA^19^, and SNP-heritability estimates were included in the modeling described above.

Tissue, cell type, and developmental time point specificity were measured using the expression specificity metric Tau.^20; 21^ Tissue specificity for each gene was calculated using publicly available expression data for 53 tissues from the GTEx project^22^ (release V6p). Expression for each tissue was summarized as the log2 of the median expression plus one, and then used to calculate tissue specificity Tau. Cell type specificity for each gene was computed using publicly available single-cell RNA-sequencing expression data^23^ generated from human cortex and hippocampus tissues. Raw expression counts for 285 cells comprising six major cell types of the brain were obtained from GEO (GSE67835) and counts data were library normalized to CPM. Expression for each cell type was then summarized as the log2 of the mean expression plus one, and then used to compute cell type specificity Tau. Developmental time point specificity for each gene was calculated using publicly available DLPFC expression data for 27 time points, clustered into eight biologically relevant groups, from the BrainSpan atlas (see Web resources). Eight developmental periods^24^ were defined as follows: early prenatal (8-12 pcw), early mid-prenatal (13-17 pcw), late mid-prenatal (19-24 pcw), late prenatal (25-37 pcw), infancy (4 mos - 1 yr), childhood (2 - 11 yr), adolescence (13 - 19 yr), and adulthood (21 yr +). Expression for each time point was summarized as the log2 of the median expression plus one, and then used to calculate developmental period specificity (Tau). Each Tau was added to the above modeling of eQTL number in turn, as well as all together.

### Enrichment Analyses

We divided eQTL into separate subgroups by stepwise conditional order (first, second, third, and greater than third), and created sets of matched SNPs drawn from the SNPsnap database for each subgroup, matching on minor allele frequency, gene density (number of genes within 1Mb of the SNP), distance from SNP to TSS of the nearest gene, and LD (number of LD-partners within r^2^ ≥ 0.8). For each subgroup of eQTL, we performed a logistic regression of status as eQTL or matched SNP on overlap with functional annotation, including the four SNP matching parameters as covariates. Enrichment was taken as the regression coefficient estimate, interpretable as the log-odds ratio for being an eQTL given a functional annotation. Functional annotations tested included: DLPFC promoters and enhancers (TssA and Enh+EnhG, respectively, from the NIH Roadmap Epigenomics Project^25^ ChromHMM^26^ core 15-state model), Brain promoters and enhancers (union of all brain region TssA and Enh+EnhG, respectively, from the NIH Roadmap Epigenomics Project ChromHMM core 15-state model), and pre-frontal cortex (PFC) neuronal (NeuN+) and non-neuronal (NeuN-) nuclei H3K4me3 and H3K27ac ChIP-seq marks from the CMC. For each data source, Roadmap DLPFC, brain, CMC NeuN+, NeuN-, active promoter and enhancer (or H3K4me3 and H3K27ac) annotation were tested for enrichment jointly.

### Conditional eQTL Analyses

In order to isolate each conditionally independent cis-eQTL association, we carried out a series of “all-but-one” conditional analyses, implemented within MatrixEQTL^15^, for each gene possessing more than one independent eQTL. As these conditional eQTL signals were to be used to test for co-localization with the SCZ GWAS signals, we limited these analyses to those genes (346 in total) with eQTL overlapping GWAS loci. For each of these genes, we conducted an ‘”all-but-one” analysis for each independent eQTL by regressing the given gene’s expression data on the dosage data, including all of the other independent eQTL for that gene as covariates in addition to diagnosis and five ancestry components. For example, three conditional analyses would be conducted for a gene with three independent eQTL: one analysis conditioning on the secondary and tertiary eQTL, one analysis conditioning on the primary and tertiary, and one analysis conditioning on the primary and secondary. In this manner we generated summary statistics for each independent eQTL in isolation, conditional on all of the other independent eQTL for that gene.

### Co-localization Analyses

For our co-localization analyses, we used summary statistics and genomic intervals from the 2014 PGC SCZ GWAS.^27^ We included 217 loci at a P-value threshold of 1x10^-6^ (omitting the complex MHC locus), defined these loci by their LD r^2^ ≥ 0.6 with the lead SNP, and then merged overlapping loci. GWAS and eQTL signatures were qualitatively compared using P-P plots, rendered in R, and LocusZoom^28^ plots.

We tested for co-localization using an updated version of COLOC^29^ R functions, which we name COLOC2 (see Web Resources) which incorporates several improvements to the method. First, COLOC2 preprocesses data by aligning eQTL and GWAS summary statistics for each eQTL cis-region. Second, the COLOC2 model optionally incorporates changes implemented in gwas-pw^30^. Briefly, we implemented learning mixture proportions of five hypotheses (*H*_*0*_, no association; *H*_*1*_, GWAS association only; *H*_*2*_, eQTL association only; *H*_*3*_, both but not co-localized; and *H*_*4*_, both and co-localized) from the data. COLOC2 uses these proportions as priors (or optionally, COLOC default or user specified priors) in the empirical Bayesian calculation of the posterior probability of co-localization for each locus (eQTL cis-region). COLOC2 averages per-SNP Wakefield asymptotic Bayes factors (WABF)^31^ across three different values for the WABF prior variance term, 0.01, 0.1, and 0.5, and provides options for specifying phenotypic variance, estimating it from case-control proportions, or estimating it from the data.

## RESULTS

### Identification of eQTL

Primary and conditional eQTL were identified using genotype and RNA-seq data from the CommonMind Consortium post-mortem DLPFC samples (467 European-ancestry cases and controls).^14^ We identified 16,273 conditional eQTL in addition to the 13,137 primary eQTL we previously reported^14^ for a total of 29,410 independent cis-eQTL for 15,817 autosomal genes. Of the genes tested, 81% (12,813 genes) had at least one eQTL and 63% of these (51% of all genes) also had at least one conditional eQTL, with an average of 1.83 independent eQTL per gene (2.26 among those with at least one eQTL), and a maximum of 16 eQTL (Figure 1). Conversely, when examining the distributions for the number of genes whose expression was affected by each eQTL (**Table S1**), the majority of eQTL were specific for a single gene, and only a small fraction of eQTL, 1.47%, affected more than one gene, with a maximum of six genes affected by a single eQTL.

**Figure 1.**
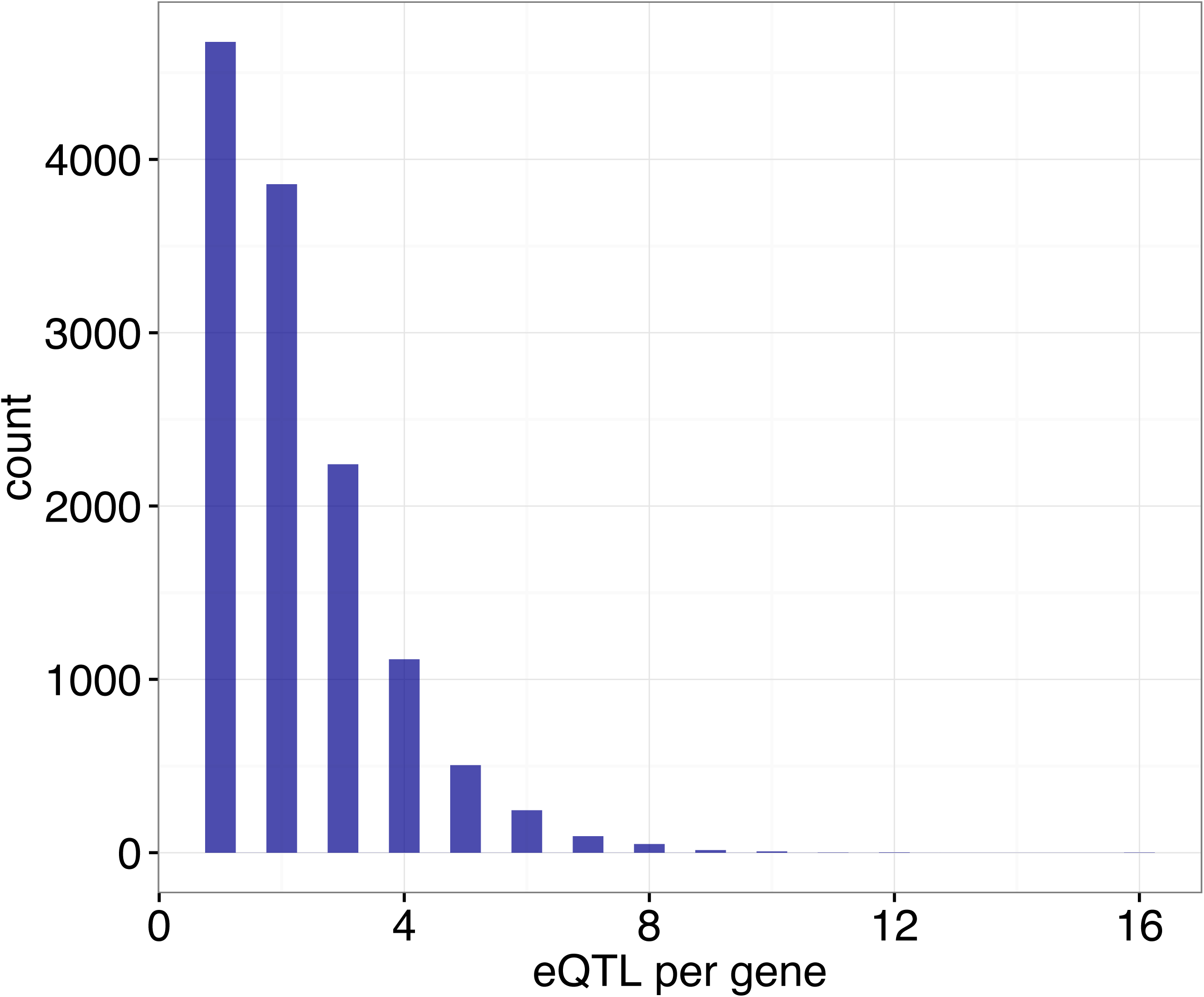
Distribution of the Number of Independent eQTL per Gene. Counts of the numbers of genes (y>axis) regulated by *N* (1 ≤ *N* ≤ 16) independent eQTL (x>axis). Plotted are 28,895 cis>eQTL with FDR ≤ 5%, for 12,813 autosomal genes. For genes with eQTL, there are an average of 2.26 eQTL per gene and a maximum of 16 eQTL per gene.

We tested conditional eQTL for replication in two independent data sets, the National Institute of Mental Health’s Human Brain Collection Core (HBCC, N=279, microarray expression data) and the Religious Orders Study / Memory and Aging Project^16^ (ROSMAP, N=494, RNA-seq expression). For each gene the same models were evaluated that were identified in forward-stepwise conditional analysis in the CMC data. We observed strong evidence of replication for both primary and conditional eQTL in the HBCC and ROSMAP post-mortem brain cohorts (**Table S2**). The estimated proportion of true associations (*p1*) in ROSMAP was 0.57 and 0.26 for primary and conditional eQTL, respectively; in HBCC *p1* was 0.46 and 0.20 for primary and conditional eQTL. Thus replication was stronger for primary than for conditional eQTL, as expected given their stronger effect sizes. Replication rates were somewhat higher in the RNA-seq ROSMAP data than in HBCC.

### Genomic Characterization of Primary and Conditional eQTL

According to prior results, eQTL that are shared across tissues and cell types tend to be located closer to transcription start sites (TSS) than context specific eQTL.^9; 12; 13^ We therefore examined the relationship between primary or conditional eQTL status and distance to its gene’s transcription start site. Primary eQTL fall closer to the TSS than conditional eQTL (Figure 2): primary eQTL occur at a median distance of 70.4 Kb from the TSS versus a median distance of 302 Kb for conditional eQTL. This difference holds true even more proximally to TSS (**Figure S1**); 8.1 and 2.5 percent of primary and conditional eQTL, respectively, fall within three Kb of the TSS.

**Figure 2.**
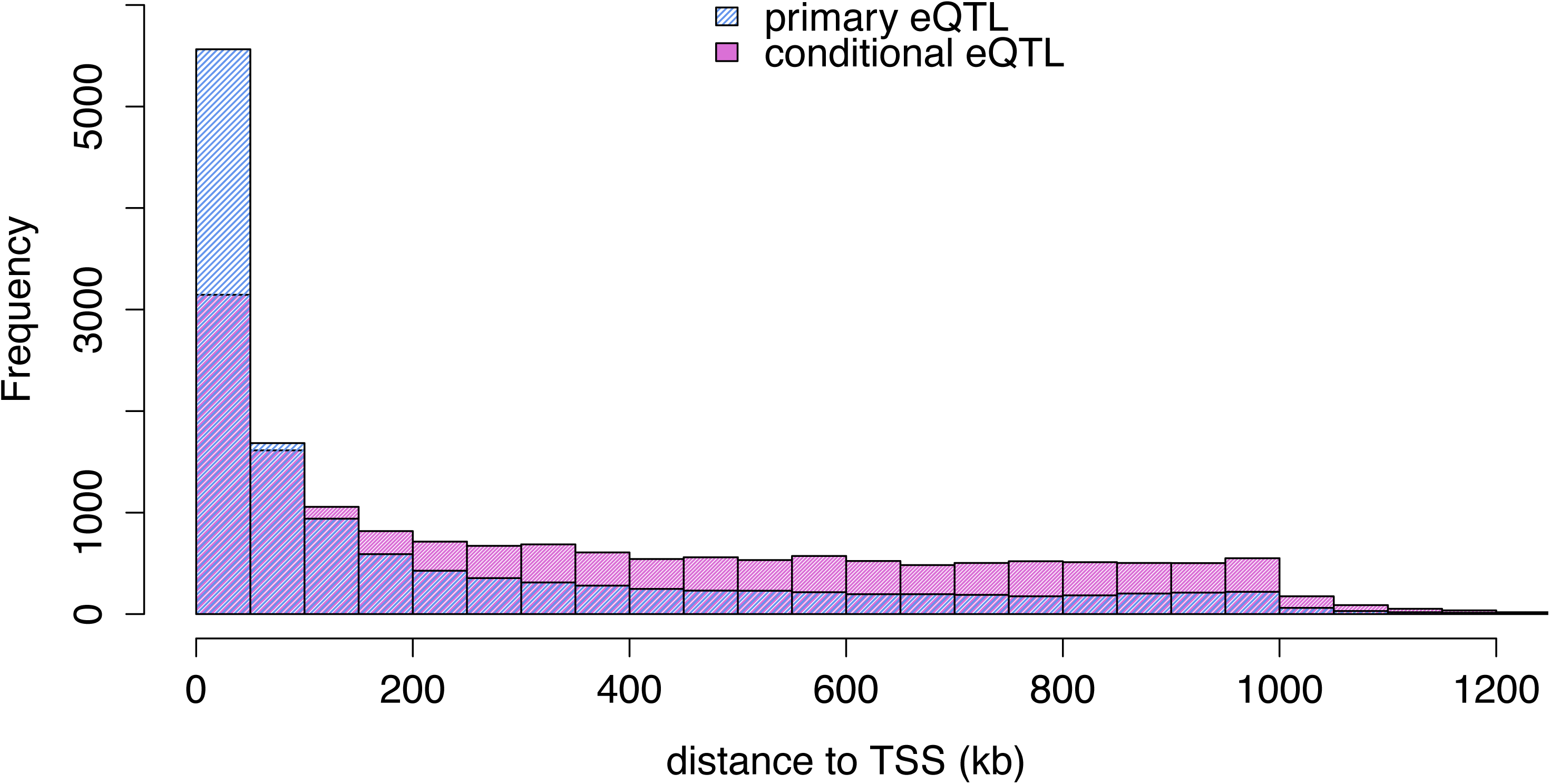
Distance from eQTL to transcription start site (TSS) Overlapping histograms showing the numbers of eQTL (yAaxis) occurring at increasing distances to TSS (xAaxis), for primary eQTL (blue) and conditional eQTL (pink).

We next characterized the relationship between the number of independent eQTL per gene and three different genomic features: gene length, number of LD blocks^32^ in the gene’s cis-region (±1 Mb) and Exome Aggregation Consortium (ExAC) genic constraint score,^33^ including possible interactions. The best multivariate model for eQTL number included gene length, number of LD blocks and genic constraint as predictors, as well as a gene length-LD blocks interaction (Table 1). The number of independent eQTL was positively correlated with gene length and number of LD blocks, and negatively correlated with genic constraint score (**Figure S2**).

**Table 1.** Number of Independent eQTL Modeled on Genomic Features.

| Predictor | Model 1<br>Estimate | Model 1<br>Robust SE | Model 1<br>Pr(>\|z\|) | Model 2<br>Estimate | Model 2<br>Robust SE | Model 2<br>Pr(>\|z\|) | Model 3<br>Estimate | Model 3<br>Robust SE | Model 3<br>Pr(>\|z\|) |
| --- | --- | --- | --- | --- | --- | --- | --- | --- | --- |
| log(Gene length) | 0.27 | 0.04 | 5.16E-12 | 0.16 | 0.03 | 2.20E-06 | 0.17 | 0.03 | 9.87E-07 |
| LD blocks | 0.59 | 0.17 | 6.47E-04 | 0.33 | 0.15 | 2.92E-02 | 0.37 | 0.15 | 1.55E-02 |
| log(Gene length) : LD<br>blocks | -0.03 | 0.02 | 7.77E-02 | -0.01 | 0.01 | 5.65E-01 | -0.01 | 0.01 | 4.11E-01 |
| Constraint | -0.61 | 0.03 | 5.93E-85 | -0.20 | 0.03 | 2.93E-13 | -0.15 | 0.03 | 5.41E-08 |
| Cis-heritability | - | - | - | 7.03 | 0.18 | 0.00 | 7.02 | 0.18 | 0.00 |
| Tau (tissue) | - | - | - | - | - | - | 0.08 | 0.08 | 2.76E-01 |
| Tau (DLPFC cell type) | - | - | - | - | - | - | 0.20 | 0.09 | 3.69E-02 |
| Tau (developmental<br>time point) | - | - | - | - | - | - | 0.17 | 0.09 | 5.99E-02 |

We next examined the variance of gene expression explained by cis-region SNPs, or cis-SNP-heritability, estimated by linear mixed model variance component analysis^19^ (**Figure S3**). We found a strong effect of estimated cis-heritability on number of independent eQTL (Table 1, **Figure S4**). In a joint model with cis-SNP-heritability, the main effects of gene length, number of LD blocks and genic constraint on eQTL number remained at least nominally significant.

Finally we addressed whether genes with conditional eQTL exhibit greater context specificity as measured by the robust expression specificity metric Tau.^20; 21^ We calculated Tau across 53 tissues from the Genotype-Tissue Expression (GTEx) project, across six DLPFC cell types (astrocytes, endothelial cells, microglia, neurons, oligodendrocytes, and oligodendrocyte progenitor cells) from single cell RNA-seq^23^, and across eight developmental periods^24^ (early prenatal, early mid-prenatal, late mid-prenatal, late prenatal, infant, child, adolescent, and adult) from the BrainSpan atlas DLPFC RNA-seq data. We confirmed that higher values of Tau reflect expression specificity, by comparing the distributions of all three Tau measures for all genes with the distributions for a subset of housekeeping genes^34^ (**Figure S5**). We found positive correlations between eQTL number and tissue, cell type, and developmental specificities (Table 1, **Table S3, Figure S6**). The strongest correlation was with DLPFC cell type Tau, which is consistent with previous data demonstrating tissue specific, cell type dependent expression in blood;^35^ however, we note that all three Tau sets were inter-correlated (**Table S3**).

### Epigenetic Enrichment Analyses

One way in which eQTL may affect gene expression is through alteration of cis-regulatory elements such as promoters and enhancers. Putative causal eQTL variants have been shown to be enriched in genomic regions containing functional annotations such as DNase hypersensitive sites, transcription factor binding sites, promoters, and enhancers.^36-39^ Our observation that conditional eQTL fall farther from transcription start sites than primary eQTL led us to hypothesize that primary eQTL may affect transcription levels by altering functional sites in promoters whereas conditional eQTL may do so by altering more distal regulatory elements such as enhancers. We therefore assessed enrichment of primary and conditional eQTL in DLPFC and brain active promoter (TssA) and enhancer (merged Enh and EnhG) states derived from the NIH Roadmap Epigenomics Project,^25; 26^ and in H3K4me3 and H3K27ac ChIP-seq peaks from a subset of the CMC post-mortem DLPFC samples. We performed logistic regression of SNP status (eQTL versus random matched SNP) on overlap with functional annotations, separately for each eQTL order (primary, secondary, tertiary, and greater than tertiary).

Both primary and conditional eQTL were significantly enriched in both promoter and enhancer chromatin states (Figure 3A-B, **Table S4**). Chromatin states from the DLPFC showed stronger eQTL enrichment than did the Brain annotation formed by merging all individual brain region chromatin states. We found that enrichments in both the DLPFC and Brain annotations generally decreased with higher conditional order of eQTL, particularly for the active promoter state. A similar pattern was observed when examining enrichment in neuronal nuclei (NeuN+) ChIP-seq peaks (Figure 3C), using the overlap of H3K4me3 and H3K27ac ChIP-seq peaks as a proxy for active promoters and H3K27ac peaks that do not overlap H3K4me3 peaks as a (relatively non-specific) proxy for enhancers.^26^ These analyses showed decreasing enrichment in both promoters and enhancers with increasing eQTL order, with a more marked decrease in the promoters. Though there was also significant enrichment of eQTL in non-neuronal nuclei (NeuN-) ChIP-seq peaks, decreasing with higher eQTL order, this trend of a more marked decrease in active promoters was not observed in non-neuronal DLPFC nuclei (Figure 3D). Enrichment results for H3K4me3 and H3K27ac ChIP-seq peaks are shown in Figure S7.

**Figure 3.**
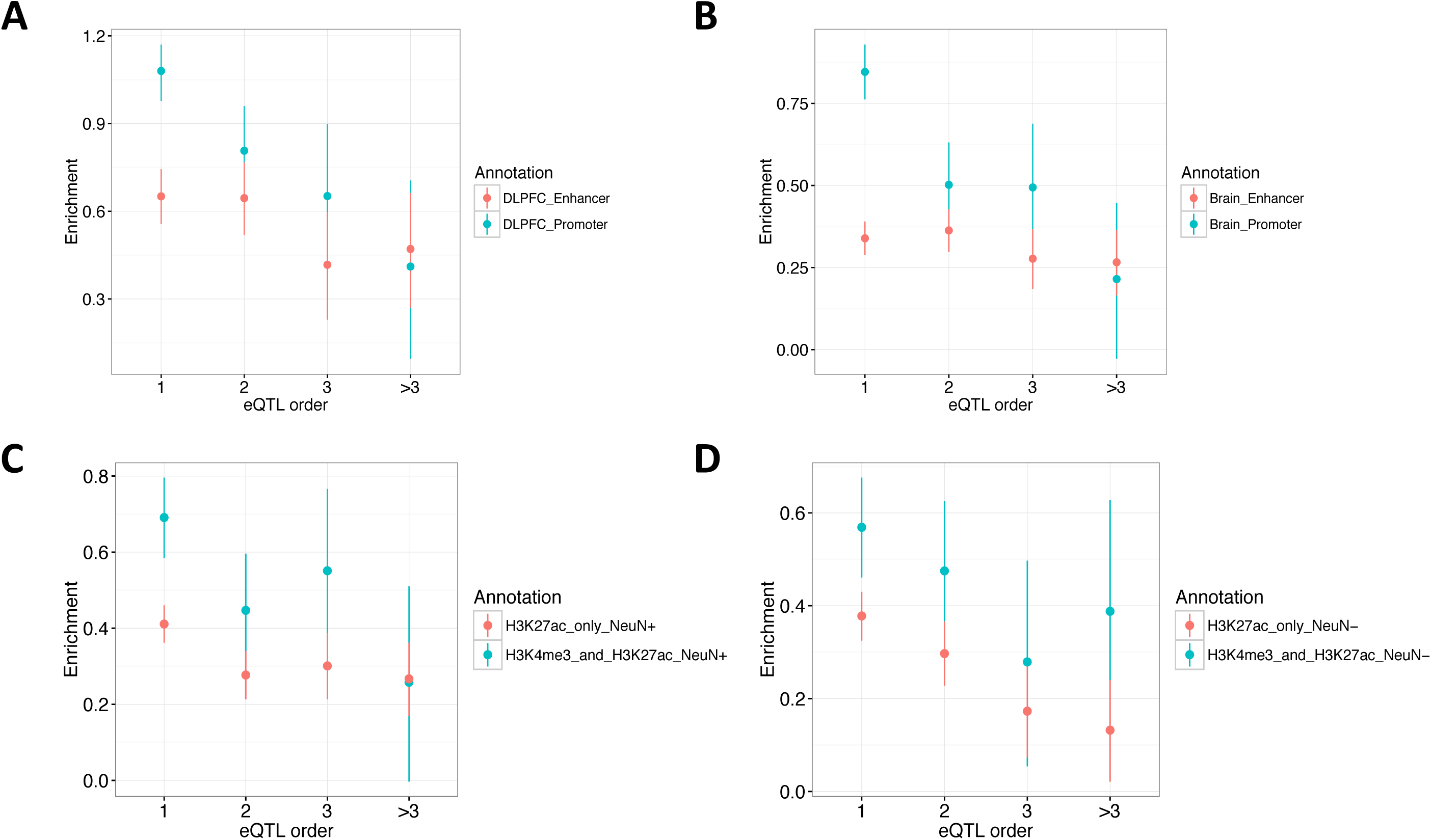
Enrichments of Primary and Conditional eQTL in Active Regulatory Annotations. Plotted are enrichments (estimate ± 95% CI from logistic regression, yFaxes) of primary (xFaxis eQTL order = 1) and conditional (eQTL order = 2, 3, >3) eQTL in functional annotations. Panels (A) and (B) show enrichment in DLPFC and Brain (union of all individual Brain regions) active promoter (turquoise) and enhancer (orange) ChromHMM states from the NIH Roadmap Epigenomics Project. Panel (C) shows enrichment in neuronal nuclei (NeuN+), for the intersection of H3K4me3 and H3K27ac ChIPFseq peaks (turquoise) and for H3K27 peaks that do not overlap H3K4me3 peaks (orange). Panel (D) shows enrichments in the same annotations, but for nonFneuronal nuclei (NeuNF).

### eQTL Co-localization with SCZ GWAS

We performed co-localization analyses in order to evaluate the extent of overlap between eQTL and GWAS signatures in schizophrenia, and to identify putative causal genes from GWAS associations. Considering 217 loci (**Table S5)** with lead SNPs reaching a significance threshold of P < 1x10^-6^ from the recent Psychiatric Genomics Consortium schizophrenia GWAS,^27^ we tabulated the number of eQTL (FDR 5%) falling within GWAS loci. A total of 114 out of 217 loci contained primary and/or conditional eQTL for 346 genes; 110 of these genes had one eQTL only, and 236 genes had more than one independent eQTL.

To quantitatively compare the SCZ GWAS and eQTL association signatures, we modified the R package COLOC^29^ for Bayesian inference of co-localization between the two sets of summary statistics across each gene’s cis-region. COLOC2, our modified implementation of COLOC, analyzes the hierarchical model of pw-GWAS,^30^ with likelihood-based estimation of dataset-wide probabilities of five hypotheses (*H*_*0*_, no association; *H*_*1*_, GWAS association only; *H*_*2*_, eQTL association only; *H*_*3*_, both but not co-localized; and *H*_*4*_, both and co-localized). We then used these probabilities as priors to calculate empirical Bayesian posterior probabilities for the five hypotheses for each locus, in particular PP_*H4*_ for co-localization.

For genes with conditional eQTL overlapping SCZ GWAS loci, summary statistics from “all-but-one” conditional eQTL analyses were assessed for co-localization with the GWAS signature (Figure 4). To illustrate this analytical strategy, we show eQTL results for the iron responsive element binding protein 2 gene *IREB2* (*chr15:78729773-78793798*) as an example. Forward stepwise selection analysis identified two independent cis-eQTL for *IREB2*. In order to generate summary statistics for each eQTL in isolation, we conducted two “all-but-one” conditional analyses, in each analysis conditioning on all but a focal independent eQTL (for *IREB2* this entailed conditioning on only one eQTL per conditional analysis, but involved conditioning on up to six eQTL across genes in the SCZ co-localization analysis). We then tested for co-localization between the GWAS and all of the conditional summary statistics using COLOC2. In the case of *IREB2*, the conditional eQTL (rs7171869) was implicated as co-localized with the GWAS signal at this locus with a posterior probability for co-localization PP_*H4*_ = 0.94. A qualitative examination of the *IREB2* locus supported the COLOC2 results: the correlation between the GWAS P-values and conditional eQTL P-values was higher than that between the GWAS and primary eQTL P-values (Figure 5A). In addition, the GWAS signature for the locus more closely resembled the conditional eQTL signature than either the non-conditional eQTL signature or the primary eQTL signature (Figure 5B).

**Figure 4.**
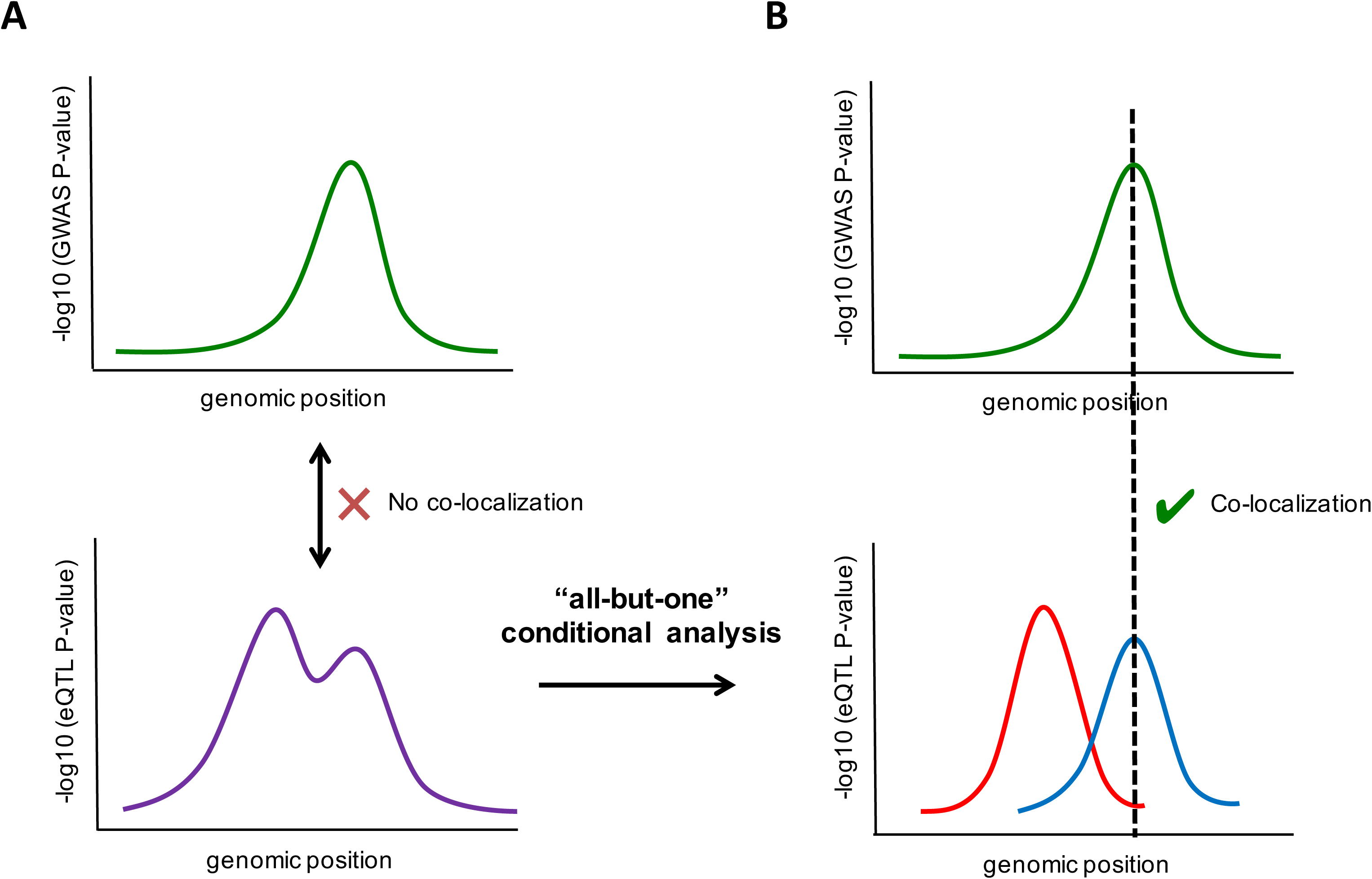
Conditional “All3but3One” Analysis to Isolate Independent eQTL Signatures. Panel (A) shows a hypothetical GWAS signature (top, green) at a given locus, and an overlapping hypothetical eQTL signature (bottom, purple), which comprises two independent eQTL. Panel (B) shows the same hypothetical GWAS and eQTL signatures after the “all3but3one” conditional eQTL analysis isolating the primary (red) and secondary (blue) eQTL signatures. Before conditional analysis there is a lack of co3 localization between the GWAS signature and eQTL signature. After all3but3one conditional analysis, there is evidence for co3localization between the secondary eQTL and GWAS signatures.

We found that 40 loci contained genes with strong evidence of co-localization between eQTL and GWAS signatures, with posterior probability of *H*_*4*_ (PP_*H4*_) ≥ 0.8 (Table 2, **Table S6**). When restricting to genome-wide significance for the GWAS, we found co-localization in 24 of the 108 loci. Given the correlations between number of independent eQTL and expression specificity scores (Tau) across tissues, cell types and development, we tabulated the reported genes’ Tau percentiles and expression levels, to highlight contexts in which the genes are specifically expressed (Table 2, **Table S9**). We acknowledge that while posterior probability PP_*H4*_ ≥ 0.8 demonstrates strong Bayesian evidence for co-localization, it is an arbitrary threshold for characterizing loci as SCZ-eQTL co-localized; we find that many loci with PP_*H4*_ ≥ 0.5 appear qualitatively consistent with co-localization.

**Table 2.** GWAS-eQTL Co-localized Loci

| Chr | Range.left | Range.right | GWAS SNP | GWAS P-value | eSNP | eSNP P-value | primary/conditional | PP<sub>H4</sub> | Gene | Relevant tissue / cell type / develop. period (See Table S9) |
| --- | --- | --- | --- | --- | --- | --- | --- | --- | --- | --- |
| 1 | 2372401 | 2402501 | rs4648845 | 4.033E-09 | rs12037821 | 6.4E-04 | conditional | 0.87 | SLC35E2 | - / - / Early mid-prenatal |
| 1 | 8355697 | 8638984 | rs301797 | 2.7E-09 | rs138050288 | 3.8E-05 | primary | 0.95 | RERE | - / - / - |
| 1 | 30412551 | 30443951 | rs1498232 | 1.3E-09 | rs2015244 | 9.7E-12 | primary | 0.99 | PTPRU | - / Neurons / Early mid-prenatal |
| 1 | 163582923 | 163766623 | rs7521492 | 5.6E-07 | rs10799961 | 2.4E-11 | primary | 0.91 | PBX1 | - / - / Early prenatal |
| 1 | 205015255 | 205189455 | rs16937 | 8.7E-07 | rs12724651 | 5.3E-07 | primary | 0.89 | TMEM81 | - / Neurons / - |
|  |  |  |  |  | rs12031350 | 8.2E-06 | conditional | 0.87 | RBBP5 | - / - / - |
| 1 | 214137889 | 214163689 | rs7529073 | 9.7E-07 | rs1431983 | 1.7E-04 | conditional | 0.93 | PROX1-AS1 | Cerebellar Hemisphere / Neurons / Adult |
| 2 | 73194203 | 73900439 | rs56145559 | 8.4E-08 | rs11679809 | 2.0E-37 | primary | 0.86 | ALMS1P | Testis / - / - |
| 2 | 110262036 | 110398236 | rs9330316 | 7.7E-08 | rs892464 | 1.3E-28 | primary | 0.92 | SEPT10 | - / - / Late prenatal |
| 2 | 198148577 | 198835577 | rs6434928 | 1.5E-11 | rs12621129 | 2.2E-12 | primary | 0.94 | SF3B1 | - / - / - |
| 2 | 200715237 | 201247789 | rs281768 | 3.5E-14 | rs35220450 | 1.6E-10 | primary | 0.95 | FTCDNL1, AC073043.2 | - / - / Adult |
|  |  |  |  |  | rs186546506 | 8.8E-04 | conditional | 0.83 | LINC01792, AC007163.3 | Putamen (basal ganglia) / - / Adult |
| 2 | 208371631 | 208531731 | rs2709410 | 4.1E-06 | rs34171849 | 2.2E-16 | primary | 0.88 | METTL21A | - / - / - |
|  |  |  |  |  | rs2551656 | 1.6E-09 | primary | 0.86 | CREB1 | - / - / Early prenatal |
| 2 | 220033801 | 220071601 | rs6707588 | 9.5E-07 | rs11236 | 1.1E-09 | primary | 0.92 | CNPPD1 | - / - / - |
| 3 | 36843183 | 36945783 | rs75968099 | 3.4E-12 | rs9834970 | 1.9E-05 | primary | 0.94 | DCLK3 | Nerve - Tibial / Neurons / Infant |
| 3 | 52281078 | 53539269 | rs2535627 | 3.956E-11 | rs6801235 | 2.8E-08 | conditional | 0.86 | PPM1M | - / Neurons / Late prenatal |
| 3 | 63792650 | 64004050 | rs832187 | 2.6E-08 | rs113386200 | 3.0E-12 | primary | 0.98 | THOC7 | - / - / - |
| 3 | 135807405 | 136615405 | rs7432375 | 5.3E-11 | rs10935184 | 7.7E-25 | primary | 0.93 | PCCB | - / - / - |
| 4 | 170357552 | 170646052 | rs10520163 | 1.0E-08 | rs7438 | 1.5E-10 | primary | 0.97 | CLCN3 | - / - / - |
| 5 | 45291475 | 46404116 | rs1501357 | 1.2E-08 | rs9292918 | 4.5E-05 | primary | 0.94 | BRCAT54, RP11-53O19.1 | - / - / Adult |
| 6 | 83779798 | 84407274 | rs3798869 | 1.2E-09 | rs2016358 | 8.7E-13 | primary | 0.90 | SNAP91 | Cerebellar Hemisphere / - / - |
| 6 | 108875527 | 109019327 | rs9398171 | 3.4E-08 | rs111727905 | 3.8E-06 | primary | 0.97 | ZNF259P1 | - / - / Early mid-prenatal |
| 7 | 21485312 | 21545712 | rs73060317 | 6.6E-07 | rs12672629 | 3.6E-05 | primary | 0.92 | SP4 | - / - / Early prenatal |
| 8 | 8088038 | 10056127 | rs2945232 | 2.0E-08 | rs2980441 | 1.9E-36 | primary | 0.82 | FAM86B3P | - / - / Adolescence |
| 8 | 26181524 | 26279124 | rs1042992 | 2.9E-07 | rs17055186 | 5.9E-25 | conditional | 0.91 | SDAD1P1 | Testis / - / Adult |
| 8 | 38020424 | 38310924 | rs57709857 | 2.3E-07 | rs17175814 | 2.1E-07 | primary | 0.88 | WHSC1L1 | - / - / Early prenatal |
| 8 | 144822546 | 144871746 | rs11784536 | 1.5E-05 | rs12541792 | 2.7E-37 | primary | 0.90 | FAM83H | Esophagus - Mucosa / Oligodendrocytes / Adolescence |
| 9 | 26839508 | 26909408 | rs10967586 | 4.7E-07 | rs12345197 | 1.3E-06 | primary | 0.80 | IFT74 | - / - / - |
| 11 | 46340213 | 46751213 | rs7951870 | 8.3E-11 | rs10160701 | 5.1E-05 | primary | 0.88 | MDK | - / - / Early mid-prenatal |
| 12 | 57428314 | 57497814 | rs324017 | 2.1E-07 | rs4559 | 4.2E-08 | conditional | 0.91 | STAT6 | - / Microglia / Adolescence |
| 14 | 35421614 | 35847614 | rs77477310 | 1.8E-07 | rs1028449 | 8.1E-04 | primary | 0.84 | RP11-85K15.2 | - / - / - |
| 15 | 78803032 | 78926732 | rs8042374 | 1.865E-12 | rs7171869 | 1.4E-04 | conditional | 0.94 | IREB2 | - / - / Early prenatal |
| 15 | 84661161 | 85153461 | rs12902973 | 8.4E-11 | rs35677834 | 1.6E-21 | primary | 0.80 | LOC101929479,<br>RP11-561C5.3 | Ovary / - / Early mid-prenatal |
| 15 | 91416560 | 91436560 | rs4702 | 2.3E-12 | rs4702 | 9.3E-10 | primary | 1.00 | FURIN | - / Endothelial cells / Adolescence |
| 16 | 4447751 | 4596451 | rs6500602 | 2.8E-07 | rs3747580 | 2.3E-16 | primary | 0.90 | CORO7 | - / - / - |
|  |  |  |  |  | rs8046295 | 4.8E-15 | primary | 0.89 | NMRAL1 | - / - / - |
| 16 | 29924377 | 30144877 | rs12691307 | 1.3E-10 | rs4788203 | 2.0E-05 | primary | 0.88 | TMEM219 | - / - / - |
|  |  |  |  |  | rs3935873 | 7.5E-14 | primary | 0.87 | INO80E | - / Neurons / - |
|  |  |  |  |  | rs4787491 | 3.5E-05 | conditional | 0.82 | DOC2A | Brain - Cortex / Neurons / Adolescence |
| 16 | 58669293 | 58691393 | rs12325245 | 1.1E-08 | rs11647976 | 4.8E-04 | primary | 0.94 | CNOT1 | - / - / - |
| 17 | 17722402 | 18030202 | rs8082590 | 6.8E-09 | rs4072739 | 3.6E-15 | primary | 0.92 | DRG2 | - / - / - |
| 19 | 11839736 | 11859736 | rs3095917 | 1.6E-06 | rs72986630 | 2.4E-07 | primary | 1.00 | ZNF823 | - / Endothelial cells / Early prenatal |
| 19 | 19374022 | 19658022 | rs2905426 | 6.9E-09 | rs2965199 | 9.2E-36 | primary | 0.87 | GATAD2A | - / - / - |
| 19 | 50067499 | 50135399 | rs56873913 | 2.2E-07 | rs5023763 | 5.5E-05 | primary | 0.93 | SNRNP70 | - / - / - |
| 22 | 41408556 | 42689414 | rs9607782 | 6.8E-12 | rs9607782 | 2.0E-04 | primary | 0.96 | RANGAP1 | - / - / - |

Importantly, for six of the 40 co-localizing loci, a conditional rather than primary eQTL co-localized with the GWAS with compelling qualitative support (Table 2, Figure 5, **Figures S8-S12**). The genes showing strong evidence for conditional eQTL co-localization include *SLC35E2*, *PROX1-AS1*, *PPM1M*, *SDAD1P1*, *STAT6*, and *IREB2*. Also notable are the occurrences of complex patterns of co-localization for some loci; for example, three loci showed evidence for co-localization with a primary eQTL for one gene and a conditional eQTL for another.

**Figure 5.**
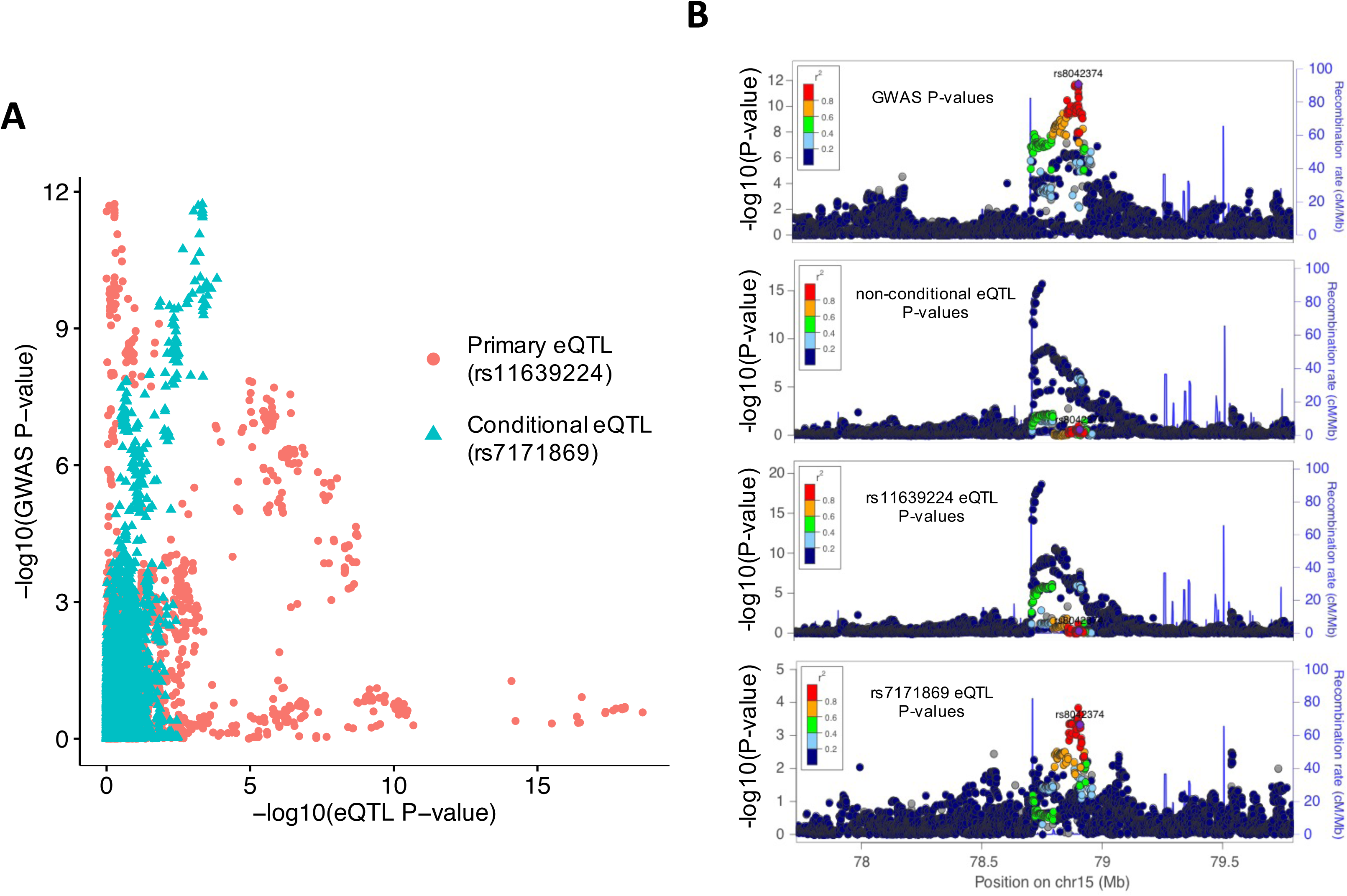
GWAS Signature for *IREB2* Co4localizes with the Conditional eQTL Signature. Panel (A) shows a P4P plot comparing 4log_10_ P4values from GWAS (y4axis) and “all4but4 one” conditional eQTL analysis (x4axis), which shows the highest correlation between the GWAS and the conditional eQTL (rs7171869, turquoise triangles). Panel (B) shows LocusZoom plots for the *IREB2* locus, where the GWAS signal (top) more closely resembles the conditional eQTL signal (rs7171869, bottom) than the primary eQTL signal (rs11639224, third from top) or non4conditional eQTL signal (second from top). For all LocusZoom plots LD is colored with respect to the GWAS lead SNP (rs8042374, labelled).

### Comparison with Previous Co-localization Analyses

In our prior analyses^14^ we reported a co-localization analysis of the 108 genome-wide significant schizophrenia GWAS loci and non-conditional eQTL using Sherlock.^40^ Those results and our current findings are highly concordant (**Table S7**). Eleven loci were reported as co-localized in both analyses. Thirteen loci were co-localized (PP_*H4*_ ≥ 0.8) in our analysis but not previously, twelve of which showed suggestive significance in Sherlock (P<2x10^-4^), or in one case involved a conditional eQTL (*SLC35E2*) in our analysis. Six loci were co-localized in the previous study but not in the current analysis; three of these resulted from differences in study design such as GWAS locus definition and eQTL overlap criteria, and two were suggestive in the current analysis (0.65< PP_*H4*_ <0.8). The one remaining discrepant locus (*chr8:143302933-143403527*) was found to co-localize with *TSNARE1* eQTL previously (Sherlock P=8.24x10^-7^), but not here (COLOC2 primary eQTL PP_*H4*_=0.074, PP_*H3*_=0.93), and indeed a qualitative comparison of the eQTL and GWAS data did not appear to support co-localization (**Figure S13**).

In the present analysis we considered not only primary but also conditional eQTL association signals for co-localization with the GWAS, allowing us to detect loci where co-localization may be obscured by multiple association signals in non-conditional eQTL analysis. We also compared our conditional co-localization results with results using non-conditional eQTL analysis, using the same COLOC2 method and SCZ GWAS loci (**Table S8**). Conditional and non-conditional COLOC2 results were highly concordant, with slightly higher PP_*H4*_s resulting from the same WABFs because of a higher prior probability of co-localization estimated in the non-conditional COLOC2 analysis. Thirty-five loci were co-localized in both analyses, and five loci that were co-localized in the non-conditional analysis only were all highly suggestive in the conditional analysis (0.65 < PP_*H4*_ < 0.8). The five loci that were co-localized only in the conditional COLOC2 analysis involved conditional and not primary eQTL.

## DISCUSSION

We utilized genotype and expression data from 467 human post-mortem brain samples from the DLPFC to conduct eQTL mapping analyses, to characterize both primary and conditionally independent eQTL. We then identified co-localization between SCZ GWAS and our eQTL association signals, including conditional eQTL. Our principal findings include four major observations. First, we detect that conditional eQTL are widespread in the brain tissue samples we investigated. In 63% of genes with at least one eQTL, we found multiple statistically independent eQTL (8,136 genes). This demonstrates that genetic variation affecting RNA abundance is incompletely characterized by focusing only on one primary eQTL per gene, which is the case currently for most eQTL studies. We suggest that these conditional eQTL may represent regulatory variation specific to biological contexts not necessarily well represented in the transcriptomic data at hand.

Second, we find the genomics of conditional eQTL and their genes are consistent with complex, context-specific regulation of gene expression. Conditional eQTL occur farther from transcription start sites than primary eQTL, consistent with effects on distal regulatory elements. Genes with more independent eQTL tend to be larger and span multiple recombination hotspot intervals, and tend to be less constrained at the protein level. While these associations may reflect in part greater power to detect independent eQTL that are not in linkage disequilibrium and that have greater phenotypic variance, they are also consistent with more complex regulation and greater potential for regulatory genetic variation. The strong association of eQTL number with gene expression cis-SNP-heritability shows that conditional eQTL contribute to regulatory genetic variation. Importantly, associations with specificity of expression across tissues, developmental periods, and cell types determined from single-cell RNA sequencing data, suggest that context specificity plays a role in the occurrence of multiple statistically independent eQTL. Cell type specificity is particularly strongly correlated with eQTL number, consistent with those cell types being present in the current tissue-homogenate data.

Both primary and conditional eQTL are enriched in both active promoter and enhancer regions, and their enrichment in active promoters diminishes with increasing conditional eQTL order. In other words, conditional eQTL show greater enrichment in enhancers relative to promoters than do primary eQTL. We note that these enrichment analyses are less well powered for conditional eQTL than for primary eQTL, both because of smaller effect sizes of conditional eQTL, and because of statistical error introduced by forward stepwise conditional analyses.^41-43^

Third, we highlight the importance of examining conditional eQTL for co-localization with GWAS. In at least six out of 40 loci showing GWAS-eQTL co-localization, a conditional eQTL signal co-localizes with SCZ risk. If we had considered only primary eQTL in the analyses, these instances of co-localization would have been missed. Conditional eQTL that co-localize with disease risk may reflect regulatory mechanisms that are important in a key developmental period or individual cell type, and may be missed when focusing on primary eQTL discovered in adult whole tissue. As further efforts are made to generate data across ranges of tissues or individual cell-types, we may have a better ability to directly identify regulatory variants specific to these contexts. However if a variant is primarily active in a very specific time point or stimulus condition, capturing data reflecting this condition will remain challenging. Conditional co-localization analysis in well-powered eQTL cohorts may best identify the genes driving these trait associations, though further validation work will be required to understand the mechanism by which the gene contributes to disease risk.

Fourth, we have identified a number of candidate genes for which genetic variation for expression co-localizes with genetic variation for schizophrenia risk (Table 2), including cases of co-localization with conditional eQTL. Genetic co-localization is expected if gene expression causally mediates disease risk, although we recognize that co-localization could also result from pleiotropy or linkage, particularly in regions of extensive linkage disequilibrium and haplotype structure.^44; 45^ Our analyses prioritize 24 genes among the 111 genes within 23 genome-wide significant SCZ loci (GWAS P < 5x10^-8^), and 22 genes in 17 suggestive (P < 1x10^-6^) loci. Among the candidates are genes that have previously been implicated in SCZ etiology, such as *FURIN*,^14^ as well as alternative candidates in well-known SCZ loci – *DCLK3* in the *TRANK1* locus,^46^ *PPM1M* in the *ITIH1* locus,^47^ *IREB2* in the *CHRNA3* locus,^27^ and *GATAD2A* in the *NCAN* locus.^27^ Our candidates include several genes not previously considered as candidates,^27^ in some cases - *SLC35E2*, *PTPRU*, *LINC01792*, *DCLK3*, *PPM1M*, *LOC101929479* - because the genes themselves do not overlap the GWAS locus regions but their eQTL do. We also find several non-coding RNA genes - *PROX1-AS1*, *FTCDNL1*, *LINC01792*, *BRCAT54*.

In an effort to highlight specific developmental periods or cell types for follow-up, we have tabulated expression specificity in GTEx tissues, brain sample cell types from single-cell RNA-seq,^23^ and in BrainSpan DLPFC developmental periods, for all identified genes (Table 2, **Table S9**). Their expression contexts show a diversity of patterns, and can provide clues to generate specific hypotheses for functional follow-up of their potential roles in SCZ. Among DLPFC cell types, we find several genes that are specific to neurons and examples of genes specific to oligodendrocytes or endothelial cells (Table 2). Among DLPFC developmental periods, we see diverse expression patterns ranging from early prenatal and broader prenatal, to perinatal (late prenatal and infancy periods), to adolescent/adult expression. We note no clear pattern of correlation between cell type and developmental expression patterns, for example neuronal cell type expressed genes include genes with prenatal, perinatal and postnatal expression. Interestingly, however, all genes broadly expressed across cell types show prenatal expression (**Table S9**).

*IREB2* (iron regulatory element binding protein 2), highlighted in our analyses as a conditional eQTL hit, is a key regulator of iron homeostasis^48; 49^ and has been implicated in neurodegenerative disorders.^50; 51^ Mouse *IREB2* homolog Irp2 knockouts exhibit impairments in coordination and balance, exploration, and nociception.^49^ The *IREB2* locus includes the *CHRNA3-CHRNA5-CHRNB4* nicotinic receptor cluster associated with schizophrenia^27^ as well as nicotine dependence and smoking behavior,^52^ lung cancer,^53; 54^ and COPD.^55^ We note that the *IREB2* conditional eQTL is associated with *CHRNA3* and *CHRNA5* expression in cerebellum, caudate and some non-brain tissues in GTEx, but both genes are too lowly expressed for eQTL analysis in the CMC DLPFC samples. Therefore, we cannot rule out the possibility that other genes may be causal of SCZ risk at this locus, perhaps in other brain regions.

A conditional eQTL for *STAT6* co-localizes with a suggestive SCZ GWAS signal (P=2x10^-7^).^27^ The immune related transcription factor STAT6 induces interleukin 4 (IL4)-mediated anti-apoptotic activity of T helper cells, and the locus is associated with migraine^56; 57^ and brain glioma,^58^ as well as several immune/inflammatory diseases.^59-61^ STAT6 also activates neuronal progenitor/stem cells and neurogenesis,^62^ making it intriguing as an immune-related SCZ candidate given recent observations about the role of complement factor 4 (*C4*) gene as a SCZ risk gene,^63^ and prior work potentially implicating microglia.^64^ Consistent with a role in immune-mediated synaptic pruning, *STAT6* expression is broadly postnatal and specific to microglia and neurons (**Table S9**).

Finally, a conditional eQTL for PROX1 Antisense RNA 1 (*PROX1-AS1*; *chr1, 214Mb*) co-localizes with a suggestive SCZ locus (P=9.7x10^-7^). The Prospero Homeobox 1 (PROX1) transcription factor, involved in development and cell differentiation in several tissues, including oligodendrocytes^65^ and GABAnergic interneurons^66^ in the brain. This lncRNA has been implicated as aberrantly expressed in several cancers, is upregulated in the cell cycle S-phase, and promotes G1/S transition in cell culture.^67^ Like *STAT6*, *PROX1-AS1* expression is specific to neurons and mature oligodendrocytes, and is expressed postnatally (**Table S9**).

In conclusion, we find that conditional eQTL are wide spread, and are consistent with complex and context specific regulation. Accounting for conditional eQTL leads to new findings of GWAS-eQTL co-localization, and generates specific hypotheses for gene expression possibly mediating disease risk.

## ACKNOWLEDGMENTS

Data were generated as part of the CommonMind Consortium supported by funding from Takeda Pharmaceuticals Company Limited, F. Hoffman-La Roche Ltd and NIH grants R01MH085542, R01MH093725, P50MH066392, P50MH080405, R01MH097276, RO1-MH-075916, P50M096891, P50MH084053S1, R37MH057881 and R37MH057881S1, HHSN271201300031C, AG02219, AG05138 and MH06692. Brain tissue for the study was obtained from the following brain bank collections: the Mount Sinai NIH Brain and Tissue Repository, the University of Pennsylvania Alzheimer’s Disease Core Center, the University of Pittsburgh NeuroBioBank and Brain and Tissue Repositories and the NIMH Human Brain Collection Core. CMC Leadership: Pamela Sklar, Joseph Buxbaum (Icahn School of Medicine at Mount Sinai), Bernie Devlin, David Lewis (University of Pittsburgh), Raquel Gur, Chang-Gyu Hahn (University of Pennsylvania), Keisuke Hirai, Hiroyoshi Toyoshiba (Takeda Pharmaceuticals Company Limited), Enrico Domenici, Laurent Essioux (F. Hoffman-La Roche Ltd), Lara Mangravite, Mette Peters (Sage Bionetworks), Thomas Lehner, Barbara Lipska (NIMH). ROSMAP study data were provided by the Rush Alzheimer’s Disease Center, Rush University Medical Center, Chicago. Data collection was supported through funding by NIA grants P30AG10161, R01AG15819, R01AG17917, R01AG30146, R01AG36836, U01AG32984, U01AG46152, the Illinois Department of Public Health, and the Translational Genomics Research Institute. The Genotype-Tissue Expression (GTEx) Project was supported by the <u>Common Fund</u> of the Office of the Director of the National Institutes of Health, and by NCI, NHGRI, NHLBI, NIDA, NIMH, and NINDS. The data used for the analyses described in this manuscript were obtained from the <u>GTEx Portal</u> on 09/05/16. BrainSpan: Atlas of the Developing Human Brain [Internet]. Funded by ARRA Awards 1RC2MH089921-01, 1RC2MH090047-01, and 1RC2MH089929-01.

## WEB RESOURCES

AMP-AD Knowledge Portal, (<u>https://www.synapse.org/ampad</u>)

BrainSpan atlas, <u>http://www.brainspan.org/</u>

CommonMind Consortium data, <u>http://www.synapse.org/CMC</u>

CommonMind Consortium ChIP-seq data, <u>https://www.synapse.org/#!Synapse:syn8040458</u>

<u>COLOC2, https://github.com/Stahl-Lab-MSSM/coloc2</u>

Darmanis et. al. single-cell RNA-seq data, <u>https://www.ncbi.nlm.nih.gov/geo/</u>, accession number GSE67835

ExAC Functional Gene Constraint, <u>http://exac.broadinstitute.org/downloads</u>

<u>GCTA, http://cnsgenomics.com/software/gcta/</u>

<u>GemTools, http://www.wpic.pitt.edu/wpiccompgen/GemTools/GemTools.htm</u>

GTEx Portal, <u>https://gtexportal.org/home/</u>

<u>HBCC microarray cohort, dbGaP (</u>ID: phs000979.v1.p1)<u>, https://www.ncbi.nlm.nih.gov/gap</u>

LDetect LD blocks, <u>https://bitbucket.org/nygcresearch/ldetect-data/overview</u>

NIH Roadmap Epigenomics Project chromatin state learning, <u>http://egg2.wustl.edu/roadmap/web_portal/chr_state_learning.html#core_15state</u>

R language and environment, <u>https://www.r-project.org/</u>

SNPsnap, <u>https://data.broadinstitute.org/mpg/snpsnap/</u>

## REFERENCES

1. Gilad, Y., Rifkin, S.A., and Pritchard, J.K. (2008). Revealing the architecture of generegulation: the promise of eQTL studies. Trends in genetics : TIG 24, 408–415.

2. Cookson, W., Liang, L., Abecasis, G., Moffatt, M., and Lathrop, M. (2009). Mapping complex disease traits with global gene expression. Nature reviews Genetics 10, 184–194.

3. Montgomery, S.B., and Dermitzakis, E.T. (2011). From expression QTLs to personalized transcriptomics. Nature reviews Genetics 12, 277–282.

4. Albert, F.W., and Kruglyak, L. (2015). The role of regulatory variation in complex traits and disease. Nature reviews Genetics 16, 197–212.

5. Moffatt, M.F., Kabesch, M., Liang, L., Dixon, A.L., Strachan, D., Heath, S., Depner, M.,von Berg, A., Bufe, A., Rietschel, E., et al. (2007). Genetic variants regulating ORMDL3 expression contribute to the risk of childhood asthma. Nature 448,470–473.

6. Speliotes, E.K., Willer, C.J., Berndt, S.I., Monda, K.L., Thorleifsson, G., Jackson, A.U.,Lango Allen, H., Lindgren, C.M., Luan, J., Magi, R., et al. (2010). Association analyses of 249,796 individuals reveal 18 new loci associated with bodymass index. Nature genetics 42, 937–948.

7. Dubois, P.C., Trynka, G., Franke, L., Hunt, K.A., Romanos, J., Curtotti, A., Zhernakova, A., Heap, G.A., Adany, R., Aromaa, A., et al. (2010). Multiple common variants for celiac disease influencing immune gene expression. Nature genetics 42, 295–302.

8. Libioulle, C., Louis, E., Hansoul, S., Sandor, C., Farnir, F., Franchimont, D., Vermeire, S., Dewit, O., de Vos, M., Dixon, A., et al. (2007). Novel Crohn disease locus identified by genome-wide association maps to a gene desert on 5p13.1 and modulates expression of PTGER4. PLoS genetics 3, e58.

9. Aguet, F., Brown, A.A., Castel, S., Davis, J.R., Mohammadi, P., Segre, A.V., Zappala, A., Abell, N.S., Fresard, L., and Gamazon, E.R. (2016). Local genetic effects on gene expression across 44 human tissues. In. (bioRxiv).

10. Jansen, R., Hottenga, J.J., Nivard, M.G., Abdellaoui, A., Laport, B., de Geus, E.J., Wright, F.A., Penninx, B.W., and Boomsma, D.I. (2017). Conditional eQTL Analysis Reveals Allelic Heterogeneity of Gene Expression. Human molecular genetics Epub ahead of print.

11. Zeng, B., Lloyd-Jones, L., Holloway, A., Marigorta, U.M., Metspalu, A., Montgomery, G.W., Esko, T., Brigham, K.L., Quyyumi, A.A., Idaghdour, Y., et al. (2016). Constraints on eQTL fine mapping in the presence of multi-site local regulation of gene expression. In. (bioRxiv).

12. Dimas, A.S., Deutsch, S., Stranger, B.E., Montgomery, S.B., Borel, C., Attar-Cohen, H., Ingle, C., Beazley, C., Gutierrez Arcelus, M., Sekowska, M., et al. (2009).Common regulatory variation impacts gene expression in a cell type-dependent manner. Science 325, 1246–1250.

13. Liu, X., Finucane, H.K., Gusev, A., Bhatia, G., Gazal, S., O'Connor, L., Bulik-Sullivan, B., Wright, F.A., Sullivan, P.F., Neale, B.M., et al. (2017). Functional Architectures of Local and Distal Regulation of Gene Expression in Multiple Human Tissues. American journal of human genetics Epub ahead of print.

14. Fromer, M., Roussos, P., Sieberts, S.K., Johnson, J.S., Kavanagh, D.H., Perumal, T.M., Ruderfer, D.M., Oh, E.C., Topol, A., Shah, H.R., et al. (2016). Gene expression elucidates functional impact of polygenic risk for schizophrenia. Nature neuroscience 19, 1442–1453.

15. Shabalin, A.A. (2012). Matrix eQTL: ultra fast eQTL analysis via large matrix operations. Bioinformatics 28, 1353–1358.

16. De Jager, P.L., Srivastava, G., Lunnon, K., Burgess, J., Schalkwyk, L.C., Yu, L., Eaton, M.L., Keenan, B.T., Ernst, J., McCabe, C., et al. (2014). Alzheimer's disease: early alterations in brain DNA methylation at ANK1, BIN1, RHBDF2 and other loci. Nature neuroscience 17, 1156–1163.

17. McCarthy, S., Das, S., Kretzschmar, W., Delaneau, O., Wood, A.R., Teumer, A., Kang, H.M., Fuchsberger, C., Danecek, P., Sharp, K., et al. (2016). A reference panel of 64,976 haplotypes for genotype imputation. Nature genetics 48, 1279–1283.

18. Das, S., Forer, L., Schonherr, S., Sidore, C., Locke, A.E., Kwong, A., Vrieze, S.I., Chew, E.Y., Levy, S., McGue, M., et al. (2016). Next-generation genotype imputation service and methods. Nature genetics 48, 1284–1287.

19. Yang, J., Lee, S.H., Goddard, M.E., and Visscher, P.M. (2011). GCTA: a tool for genome-wide complex trait analysis. American journal of human genetics 88, 76–82.

20. Yanai, I., Benjamin, H., Shmoish, M., Chalifa-Caspi, V., Shklar, M., Ophir, R., Bar-Even, A., Horn-Saban, S., Safran, M., Domany, E., et al. (2005). Genome-wide midrange transcription profiles reveal expression level relationships in human tissue specification. Bioinformatics 21, 650–659.

21. Kryuchkova-Mostacci, N., and Robinson-Rechavi, M. (2016). A benchmark of 760 gene expression tissue-specificity metrics. Briefings in bioinformatics 18,205–214.

22. Consortium, G. (2013). The Genotype-Tissue Expression (GTEx) project. Nature genetics 45, 580–585.

23. Darmanis, S., Sloan, S.A., Zhang, Y., Enge, M., Caneda, C., Shuer, L.M., Hayden Gephart, M.G., Barres, B.A., and Quake, S.R. (2015). A survey of human brain transcriptome diversity at the single cell level. Proc Natl Acad Sci U S A 112, 7285–7290.

24. Lin, G.N., Corominas, R., Lemmens, I., Yang, X., Tavernier, J., Hill, D.E., Vidal, M., Sebat, J., and Iakoucheva, L.M. (2015). Spatiotemporal 16p11.2 protein network implicates cortical late mid-fetal brain development and KCTD13-Cul3-RhoA pathway in psychiatric diseases. Neuron 85, 742–754.

25. Roadmap Epigenomics, C., Kundaje, A., Meuleman, W., Ernst, J., Bilenky, M., Yen, A., Heravi-Moussavi, A., Kheradpour, P., Zhang, Z., Wang, J., et al. (2015). Integrative analysis of 111 reference human epigenomes. Nature 518, 317–330.

26. Ernst, J., and Kellis, M. (2012). ChromHMM: automating chromatin-state discovery and characterization. Nat Methods 9, 215–216.

27. Schizophrenia Working Group of the Psychiatric Genomics, C. (2014). Biological insights from 108 schizophrenia-associated genetic loci. Nature 511, 421–427.

28. Pruim, R.J., Welch, R.P., Sanna, S., Teslovich, T.M., Chines, P.S., Gliedt, T.P., Boehnke, M., Abecasis, G.R., and Willer, C.J. (2010). LocusZoom: regional visualization of genome-wide association scan results. Bioinformatics 26,2336–2337.

29. Giambartolomei, C., Vukcevic, D., Schadt, E.E., Franke, L., Hingorani, A.D., Wallace, C., and Plagnol, V. (2014). Bayesian test for colocalisation between pairs of genetic association studies using summary statistics. PLoS genetics 10, e1004383.

30. Pickrell, J.K., Berisa, T., Liu, J.Z., Segurel, L., Tung, J.Y., and Hinds, D.A. (2016). Detection and interpretation of shared genetic influences on 42 human traits. Nature genetics 48, 709–717.

31. Wakefield, J. (2009). Bayes factors for genome-wide association studies: comparison with P-values. Genet Epidemiol 33, 79–86.

32. Berisa, T., and Pickrell, J.K. (2016). Approximately independent linkage disequilibrium blocks in human populations. Bioinformatics 32, 283–285.

33. Lek, M., Karczewski, K.J., Minikel, E.V., Samocha, K.E., Banks, E., Fennell, T., O'Donnell-Luria, A.H., Ware, J.S., Hill, A.J., Cummings, B.B., et al. (2016). Analysis of protein-coding genetic variation in 60,706 humans. Nature 536, 285–291.

34. Eisenberg, E., and Levanon, E.Y. (2013). Human housekeeping genes, revisited. Trends in genetics : TIG 29, 569–574.

35. Zhernakova, D.V., Deelen, P., Vermaat, M., van Iterson, M., van Galen, M., Arindrarto, W., van 't Hof, P., Mei, H., van Dijk, F., Westra, H.J., et al. (2017). Identification of context-dependent expression quantitative trait loci in whole blood. Nature genetics 49, 139–145.

36. Farh, K.K., Marson, A., Zhu, J., Kleinewietfeld, M., Housley, W.J., Beik, S., Shoresh, N., Whitton, H., Ryan, R.J., Shishkin, A.A., et al. (2015). Genetic and epigeneticfine mapping of causal autoimmune disease variants. Nature 518, 337–343.

37. Tak, Y.G., and Farnham, P.J. (2015). Making sense of GWAS: using epigenomicsand genome engineering to understand the functional relevance of SNPs innon-coding regions of the human genome. Epigenetics Chromatin 8, 57.

38. Brown, C.D., Mangravite, L.M., and Engelhardt, B.E. (2013). Integrative modelingof eQTLs and cis-regulatory elements suggests mechanisms underlying celltype specificity of eQTLs. PLoS genetics 9, e1003649.

39. Gaffney, D.J., Veyrieras, J.B., Degner, J.F., Pique-Regi, R., Pai, A.A., Crawford, G.E.,Stephens, M., Gilad, Y., and Pritchard, J.K. (2012). Dissecting the regulatoryarchitecture of gene expression QTLs. Genome Biol 13, R7.

40. He, X., Fuller, C.K., Song, Y., Meng, Q., Zhang, B., Yang, X., and Li, H. (2013). Sherlock: detecting gene-disease associations by matching patterns of expression QTL and GWAS. American journal of human genetics 92, 667–680.

41. Servin, B., and Stephens, M. (2007). Imputation-based analysis of associationstudies: candidate regions and quantitative traits. PLoS genetics 3, e114.

42. Spain, S.L., and Barrett, J.C. (2015). Strategies for fine-mapping complex traits. Human molecular genetics 24, R111–119.

43. Hormozdiari, F., Kostem, E., Kang, E.Y., Pasaniuc, B., and Eskin, E. (2014).Identifying causal variants at loci with multiple signals of association. Genetics 198, 497–508.

44. Guo, H., Fortune, M.D., Burren, O.S., Schofield, E., Todd, J.A., and Wallace, C.(2015). Integration of disease association and eQTL data using a Bayesiancolocalisation approach highlights six candidate causal genes in immune-mediated diseases. Human molecular genetics 24, 3305–3313.

45. Zhu, Z., Zhang, F., Hu, H., Bakshi, A., Robinson, M.R., Powell, J.E., Montgomery, G.W., Goddard, M.E., Wray, N.R., Visscher, P.M., et al. (2016). Integration ofsummary data from GWAS and eQTL studies predicts complex trait genetargets. Nature genetics 48, 481–487.

46. Schizophrenia Psychiatric Genome-Wide Association Study, C. (2011). Genome-wide association study identifies five new schizophrenia loci. Nature genetics 43, 969–976.

47. Sleiman, P., Wang, D., Glessner, J., Hadley, D., Gur, R.E., Cohen, N., Li, Q., Hakonarson, H., and Janssen, C.N.G.W.G. (2013). GWAS meta analysis identifies TSNARE1 as a novel Schizophrenia / Bipolar susceptibility locus. Scientific reports 3, 3075.

48. Kim, W.J., and Lee, S.D. (2015). Candidate genes for COPD: current evidence and 844 research. International journal of chronic obstructive pulmonary disease 10, 845 2249–2255.

49. Zumbrennen-Bullough, K.B., Becker, L., Garrett, L., Holter, S.M., Calzada-Wack, J., Mossbrugger, I., Quintanilla-Fend, L., Racz, I., Rathkolb, B., Klopstock, T., et al.(2014). Abnormal brain iron metabolism in Irp2 deficient mice is associatedwith mild neurological and behavioral impairments. PLoS one 9, e98072.

50. Dusek, P., Jankovic, J., and Le, W. (2012). Iron dysregulation in movement disorders. Neurobiol Dis 46, 1–18.

51. Rouault, T.A. (2013). Iron metabolism in the CNS: implications for neurodegenerative diseases. Nat Rev Neurosci 14, 551–564.

52. Thorgeirsson, T.E., Geller, F., Sulem, P., Rafnar, T., Wiste, A., Magnusson, K.P., Manolescu, A., Thorleifsson, G., Stefansson, H., Ingason, A., et al. (2008). A variant associated with nicotine dependence, lung cancer and peripheral arterial disease. Nature 452, 638–642.

53. Amos, C.I., Wu, X., Broderick, P., Gorlov, I.P., Gu, J., Eisen, T., Dong, Q., Zhang, Q., Gu, X., Vijayakrishnan, J., et al. (2008). Genome-wide association scan of tag SNPs identifies a susceptibility locus for lung cancer at 15q25.1. Nature genetics 40, 616–622.

54. Hung, R.J., McKay, J.D., Gaborieau, V., Boffetta, P., Hashibe, M., Zaridze, D., Mukeria, A., Szeszenia-Dabrowska, N., Lissowska, J., Rudnai, P., et al. (2008). A susceptibility locus for lung cancer maps to nicotinic acetylcholine receptor subunit genes on 15q25. Nature 452, 633–637.

55. DeMeo, D.L., Mariani, T., Bhattacharya, S., Srisuma, S., Lange, C., Litonjua, A., Bueno, R., Pillai, S.G., Lomas, D.A., Sparrow, D., et al. (2009). Integration of genomic and genetic approaches implicates IREB2 as a COPD susceptibility gene. American journal of human genetics 85, 493–502.

56. Anttila, V., Winsvold, B.S., Gormley, P., Kurth, T., Bettella, F., McMahon, G., Kallela, M., Malik, R., de Vries, B., Terwindt, G., et al. (2013). Genome-wide meta-analysis identifies new susceptibility loci for migraine. Nature genetics 45, 912–917.

57. Gormley, P., Anttila, V., Winsvold, B.S., Palta, P., Esko, T., Pers, T.H., Farh, K.H., Cuenca-Leon, E., Muona, M., Furlotte, N.A., et al. (2016). Meta-analysis of 375,000 individuals identifies 38 susceptibility loci for migraine. Nature genetics 48, 856–866.

58. Ruan, Z., Zhao, Y., Yan, L., Chen, H., Fan, W., Chen, J., Wu, Q., Qian, J., Zhang, T., Zhou, K., et al. (2011). Single nucleotide polymorphisms in IL-4Ra, IL-13 and STAT6 genes occurs in brain glioma. Front Biosci (Elite Ed) 3, 33–45.

59. Sleiman, P.M., Wang, M.L., Cianferoni, A., Aceves, S., Gonsalves, N., Nadeau, K., Bredenoord, A.J., Furuta, G.T., Spergel, J.M., and Hakonarson, H. (2014). GWAS identifies four novel eosinophilic esophagitis loci. Nat Commun 5, 5593.

60. Granada, M., Wilk, J.B., Tuzova, M., Strachan, D.P., Weidinger, S., Albrecht, E., Gieger, C., Heinrich, J., Himes, B.E., Hunninghake, G.M., et al. (2012). A genome-wide association study of plasma total IgE concentrations in the Framingham Heart Study. J Allergy Clin Immunol 129, 840–845 e821.

61. Bonnelykke, K., Matheson, M.C., Pers, T.H., Granell, R., Strachan, D.P., Alves, A.C., Linneberg, A., Curtin, J.A., Warrington, N.M., Standl, M., et al. (2013). Meta-analysis of genome-wide association studies identifies ten loci influencing allergic sensitization. Nature genetics 45, 902–906.

62. Bhattarai, P., Thomas, A.K., Cosacak, M.I., Papadimitriou, C., Mashkaryan, V., Froc, C., Reinhardt, S., Kurth, T., Dahl, A., Zhang, Y., et al. (2016). IL4/STAT6 Signaling Activates Neural Stem Cell Proliferation and Neurogenesis upon Amyloid-beta42 Aggregation in Adult Zebrafish Brain. Cell Rep 17, 941–948.

63. Sekar, A., Bialas, A.R., de Rivera, H., Davis, A., Hammond, T.R., Kamitaki, N., Tooley, K., Presumey, J., Baum, M., Van Doren, V., et al. (2016). Schizophrenia risk from complex variation of complement component 4. Nature 530, 177–183.

64. Schafer, D.P., Lehrman, E.K., Kautzman, A.G., Koyama, R., Mardinly, A.R., Yamasaki, R., Ransohoff, R.M., Greenberg, M.E., Barres, B.A., and Stevens, B. (2012). Microglia sculpt postnatal neural circuits in an activity and complement-dependent manner. Neuron 74, 691–705.

65. Kato, K., Konno, D., Berry, M., Matsuzaki, F., Logan, A., and Hidalgo, A. (2015). Prox1 Inhibits Proliferation and Is Required for Differentiation of the Oligodendrocyte Cell Lineage in the Mouse. PLoS one 10, e0145334.

66. Miyoshi, G., Young, A., Petros, T., Karayannis, T., McKenzie Chang, M., Lavado, A., Iwano, T., Nakajima, M., Taniguchi, H., Huang, Z.J., et al. (2015). Prox1 Regulates the Subtype-Specific Development of Caudal Ganglionic Eminence-Derived GABAergic Cortical Interneurons. J Neurosci 35, 12869–12889.

67. Yang, F., Yi, F., Zheng, Z., Ling, Z., Ding, J., Guo, J., Mao, W., Wang, X., Wang, X., Ding, X., et al. (2012). Characterization of a carcinogenesis-associated long non-coding RNA. RNA Biol 9, 110–116.

